# Downregulated NPAS4 in multiple brain regions is associated with Major Depressive Disorder

**DOI:** 10.1101/2022.08.23.505036

**Authors:** Berkay Selçuk, Tuana Aksu, Onur Dereli, Ogun Adebali

## Abstract

Major Depressive Disorder (MDD) is a commonly observed psychiatric disorder that affects more than 2% of the world population with a rising trend. However, disease-associated pathways and biomarkers are yet to be fully comprehended. In this study, we analyzed previously generated RNA-seq data across seven different brain regions from three distinct studies to identify differentially and co-expressed genes for patients with MDD. Differential gene expression (DGE) analysis revealed that NPAS4 is the only gene downregulated in three different brain regions. Furthermore, co-expressing gene modules responsible for glutamatergic signaling are negatively enriched in these regions. We used the results of both DGE and co-expression analyses to construct a novel MDD-associated pathway. In our model, we propose that disruption in glutamatergic signaling-related pathways might be associated with the downregulation of NPAS4 and many other immediate-early genes (IEGs) that control synaptic plasticity. In addition to DGE analysis, we identified the relative importance of KEGG pathways in discriminating MDD phenotype using a machine learning-based approach. We anticipate that our study will open doors to developing better therapeutic approaches targeting glutamatergic receptors in the treatment of MDD.

## Introduction

Major Depressive Disorder (MDD), also known as depression, is a common psychiatric disorder that affected more than 2% of the world population (163 million people) in 2017 (James et al., 2018). It is characterized by low mood sustained for at least 2 weeks, often with low self-esteem, loss of interest in normally enjoyable activities, low energy, and pain without a clear cause. Among more severe symptoms, suicidal behaviors are observed in patients with major depression, making it one of the most common fatal disorders in the world (National Institute of Mental Health, 2021). Recently, the severe depression rate among youth escalated from 9.4% to 21.1% between 2013 and 2018 (Duffy et al., 2019). This suggests a rising trend in the number of depressive patients and emphasizes the importance and urgency of the problem. Therefore, immediate research is needed to define fine-established markers of major depression to address this ongoing global well-being problem.

Several attempts have been made to identify the transcriptional profiles of patients with major depression by using next-generation sequencing (NGS) data obtained from post-mortem patients. Pantazatos et al. (2017) have discovered thirty-five differentially expressed genes in the dorsolateral prefrontal cortex of depression sudden deaths (MDD) and depression suicidals (MDD-S) compared to the control group (padj < 0.1). However, only the dorsolateral prefrontal cortex, with a limited sample size of 59, was investigated in that study. Labonté et al. (2017) examined six brain regions and showed differences in transcriptional patterns of men and women, proposing sexual dimorphism for depression. Although researchers have discovered a 5-10% overlap for the differentially expressed genes for the females and males, the data did not yield any outstanding common genetic marker associated with MDD. Similarly, in 2017, Ramaker et al. (2017) investigated transcriptional profiles of patients with schizophrenia, bipolar disorder, and major depression. Although they have identified differentially expressed genes (padj < 0.05) for schizophrenia and bipolar disorder, they have not identified any for major depression. Sequencing data from these three valuable studies can be analyzed together to increase the sample size and improve the resolution of the results.

In this study, we combined and analyzed previously used RNA-seq data from multiple studies (Labonté et al., 2017; Pantazatos et al., 2017; Ramaker et al., 2017) to identify genes that are differentially expressed for MDD by considering the factors of gender, age, postmortem interval, brain region, and the study they belonged to. We investigated genes that are differentially expressed in 7 distinct brain regions, including the dorsolateral prefrontal cortex (DLPFC), nucleus accumbens (nACC), ventral subiculum (vSUB), anterior insula (aINS), anterior cingulate cortex (AnCg), cingulate gyrus 25 (Cg25), and orbitofrontal cortex (OFC). Three of these brain regions (DLPFC, nACC, and vSUB) were further studied for significant gene expression changes and co-expressing gene modules. Lastly, used a non-linear, machine learning based approach to determine biological pathways that can be used for diagnostic purposes. We present significant genetic biomarkers and pathways associated with the major depression phenotype.

## Results

We combined RNA-seq datasets from three different sources (Labonté et al., 2017; Pantazatos et al., 2017; Ramaker et al., 2017) containing sequenced brain tissue samples from post-mortem control and major depression patients to identify statistically significant transcriptional changes. We analyzed the raw RNA sequencing reads and measured the expression levels of genes for each sample. The quality of each sample was assessed, and a few samples were discarded from the analysis due to having low quality (see methods). Then, we followed the general pipeline of RNA-seq data analysis (see methods) by performing alignment to the human genome and counting the reads aligned with each gene. We grouped the counts according to the brain region they belonged to and identified genes that are differentially expressed relative to the control group (padj < 0.05) for each region by using the DESeq2 R package (Love et al., 2014). We did not apply any log-fold change cut-off to our analysis.

To investigate the potential transcriptional similarities between different brain regions, we first calculated pairwise Spearman’s correlations using log2FC values of commonly expressed genes **(Figure 1A)**. No strong correlation was observed between the two regions. The strongest correlation was observed between the orbitofrontal cortex and ventral subiculum with pairwise Spearman’s correlation score of 0.28. Therefore, we can conclude that different disease-related signatures were observed in different brain regions.

**Figure 1.**
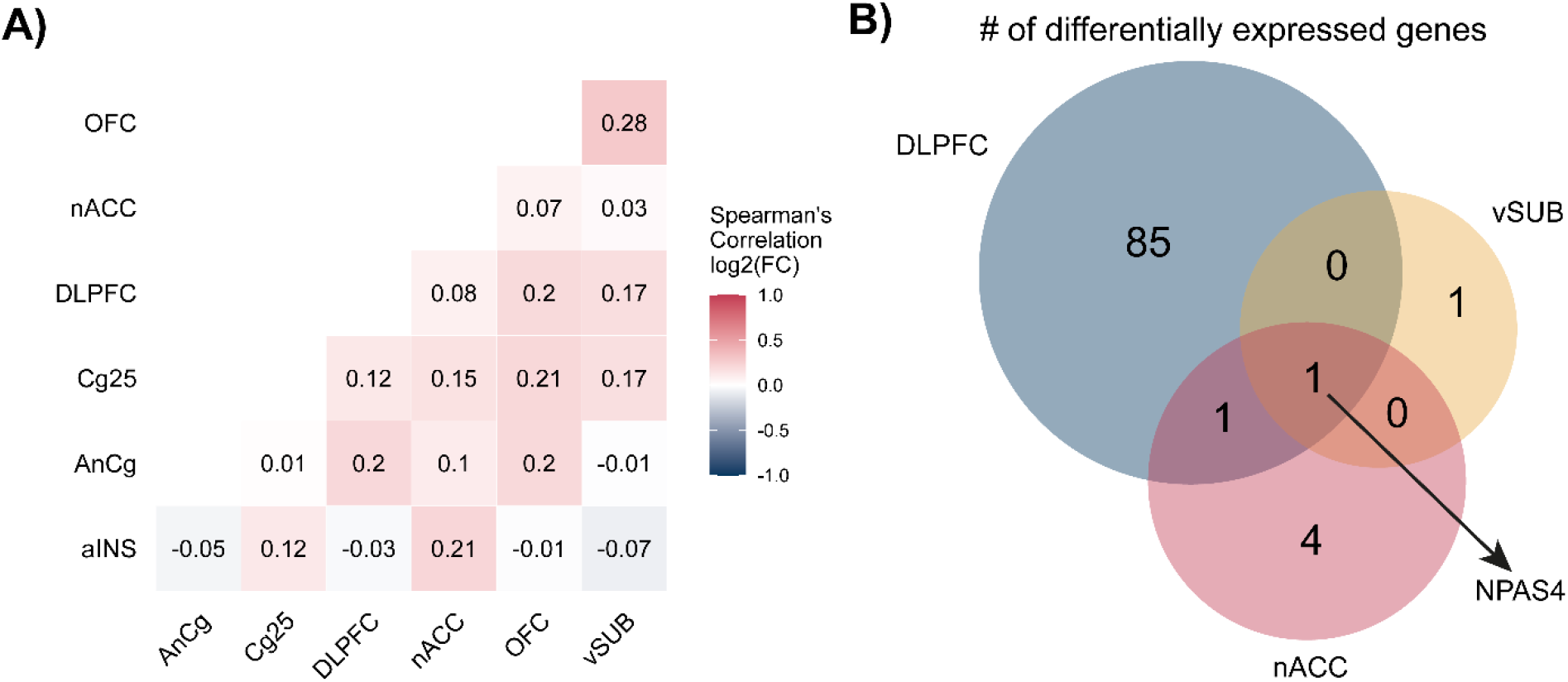
Differential gene expression analysis for different brain regions A) Spearman’s correlation of log2FC values between investigated brain regions. B) Venn diagram showing the number of differentially expressed genes for DLPC, vSUB, and nACC.

Then, we focused on genes that are differentially expressed for each region independently. Out of the seven regions, we identified at least one differentially expressed gene in DLPFC, nACC, and vSUB but not in other brain regions. The highest number of differentially expressed genes was observed in DLPFC (sample size of n=150) with 87 differentially expressed genes, and this was followed by nACC (n=94) with six genes and vSUB (n=43) with two genes **(Figure 1B)**. When we intersected lists of differentially expressed genes for these three regions, we discovered that a brain-specific transcription factor NPAS4 (Greb-Markiewicz et al., 2018) was the only common gene (Figure 1b) that was downregulated in all three regions. It was previously shown in mice (Coutellier et al., 2012; Coutellier et al., 2015; Jaehne et al., 2015; Wang et al., 2019) and in a study monitoring 152 ischemic stroke patients (Gu et al., 2019) that decrease in NPAS4 expression is correlated with the MDD phenotype.

Because NPAS4 was identified as the single common downregulated gene, we aimed to further investigate the shared transcriptional profile between different regions. Therefore, we combined samples from three regions (DLPFC, nACC, and vSUB) which we observed differential gene expression and reached a sample size of 287 (143 CTRL, 144 MDD) to perform a DGE analysis by adding a covariate of “brain region” to eliminate region-specific variations in gene expression. As presented in the volcano plot **(Figure 2a)**, 149 genes were found to be differentially expressed (padj<0.05) with a general trend of downregulation. We suggest that this was mainly due to the top three (padj:2×10^−27^, 4.9×10^−11^, 2.3×10^−8^) downregulated transcription factors (NPAS4, FOS, and FOSB) **(Figure 2b)**.

**Figure 2.**
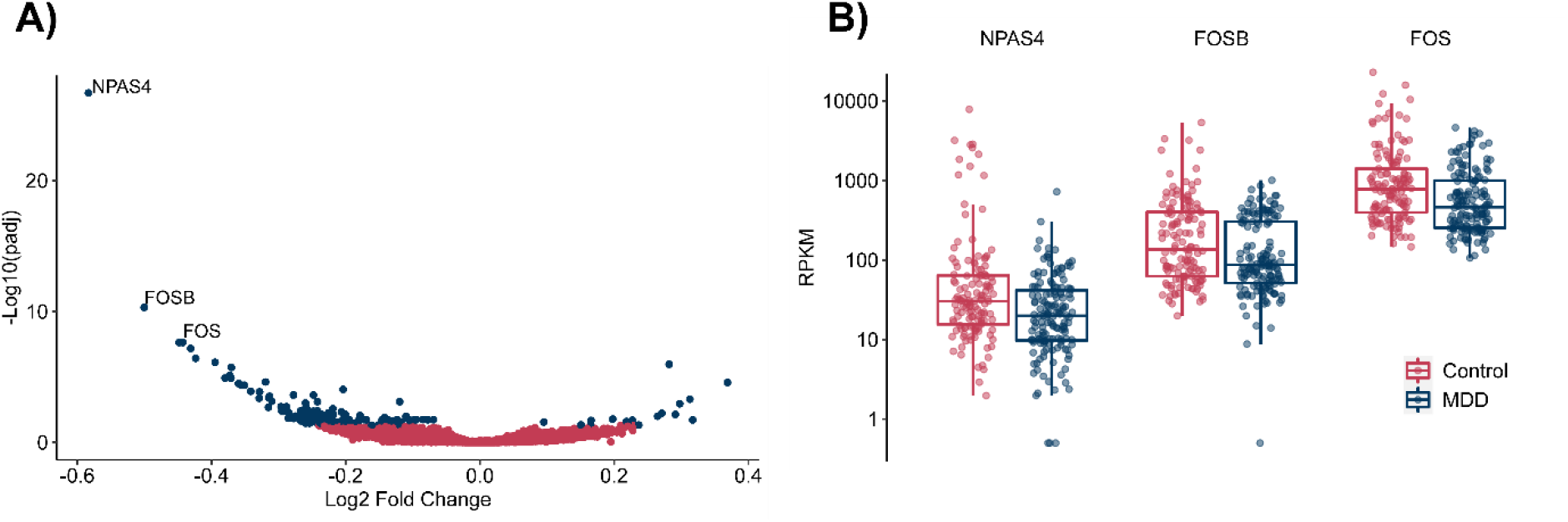
Volcano plot and top 3 genes. A) Volcano plot for the DGE analysis of three regions. Blue: padj<0.05, Red: padj ≥ 0.05 B) Box plots for the top three differentially expressed genes NPAS4, FOSB, and FOS.

To gain more insight into the pathways involved in MDD phenotype, we performed a co-expression analysis using CEMiTool (Russo et al., 2018) for the brain regions we observed NPAS4 downregulation to reveal correlating gene modules. As an input, we used the same normalized count matrix for DGE analysis. The co-expression analysis yielded two co-expressed gene modules (padj < 0.1) as modules 1 and 2. After introducing sample annotations as MDD and control, we identified that both of the modules show positive enrichment in control patients and negative enrichment in MDD patients (Supplementary Figure 1). For the first module (128 genes) control group had normalized enrichment score (NES) of 1.49 (padj = 0.048) and MDD group had −1.48 (padj=0.036). Furthermore, for the second gene (60 genes) module control group had NES of 1.42 and MDD group had −1.44 (padj=0.065). Overall, higher enrichment means a higher activity of the module for a given group and the opposite is true for the negatively enriched group as well. Because the activity of each module is correlated with the expression levels of the samples, we can conclude that they are downregulated for patients with depression. Although we have also performed this analysis by including all available samples, we did not obtain any meaningful functional enrichment for the identified modules. We explored the functional implications of the modules in the following paragraphs.

We performed gene set enrichment analysis through a web-based tool Enrichr (Chen et al., 2013; Kuleshov et al., 2016; Xie et al., 2021), and presented the top 10 KEGG (Kyoto Encyclopedia of Genes and Genomes) (Kanehisa, 2019; Kanehisa et al., 2020) pathways based on their combined score (**Table 1-3**) for differentially genes and co-expressed gene modules. Enrichment of differentially expressed genes yielded 18 pathways (padj<0.05) related to inflammation, such as the IL17 signaling pathway, Rheumatoid arthritis, NF-kappa B signaling pathway. It has been previously suggested that IL-17A induces depressive behavior in mice (Kim et al., 2021; Nadeem et al., 2017), but studies for humans (Saraykar et al., 2017; Tsuboi et al., 2018; Zafiriou et al., 2021) have contradicting conclusions. Lui et al. (2011) showed that higher serum levels of IL-17 are positively correlated with the severity of anxiety in patients with rheumatoid arthritis. The involvement of interleukins and cytokines was previously discussed numerous times (Dowlati et al., 2010; Himmerich et al., 2019; Schiepers et al., 2005). It should be noted that **91** out of **147** differentially expressed genes (including NPAS4) are not present in any KEGG pathway. This suggests that these pathways should not be considered representatives of all differentially expressed genes.

**Table 1.** KEGG 2021 pathway enrichment for the differentially expressed genes. Overlap: The overlap between the gene set and the pathway.

| *KEGG Pathway* | *Overlap* | *p<sub>adj</sub>* | *Odds Ratio* | *Combined Score* |
| --- | --- | --- | --- | --- |
| IL-17 signaling pathway | 9/94 | 5.20E-06 | 15.17 | 262.43 |
| TNF signaling pathway | 8/112 | 1.49E-04 | 10.93 | 144.84 |
| Rheumatoid arthritis | 7/93 | 3.18E-04 | 11.49 | 138.95 |
| Legionellosis | 5/57 | 0.001585773 | 13.41 | 129.97 |
| Bladder cancer | 4/41 | 0.0048795 | 14.98 | 125.53 |
| AGE-RAGE signaling pathway in diabetic | 7/100 | 3.86E-04 | 10.62 | 123.33 |
| NF-kappa B signaling pathway | 7/104 | 4.00E-04 | 10.18 | 115.59 |
| Malaria | 4/50 | 0.008858662 | 12.04 | 91.64 |
| MAPK signaling pathway | 10/294 | 0.001585773 | 5.03 | 48.47 |
| Kaposi sarcoma-associated herpesvirus infection | 7/193 | 0.008858662 | 5.29 | 39.48 |

**Table 2.** KEGG pathway enrichment for the glutamatergic signaling co-expression module (Module 1).

| *KEGG Pathway* | *Overlap* | *p**adj* | *Odds Ratio* | *Combined Score* |
| --- | --- | --- | --- | --- |
| Amphetamine addiction | 8/69 | 1.04E-06 | 22.02 | 401.59 |
| Cocaine addiction | 6/49 | 1.79E-05 | 23.06 | 329.55 |
| Circadian entrainment | 9/97 | 1.04E-06 | 17.30 | 317.75 |
| Glutamatergic synapse | 9/114 | 2.51E-06 | 14.48 | 245.48 |
| Gastric acid secretion | 7/76 | 1.76E-05 | 16.88 | 244.71 |
| Long-term potentiation | 6/67 | 9.99E-05 | 16.24 | 201.68 |
| Dopaminergic synapse | 9/132 | 6.70E-06 | 12.35 | 193.68 |
| GABAergic synapse | 6/89 | 3.31E-04 | 11.92 | 128.40 |
| Morphine addiction | 6/91 | 3.44E-04 | 11.64 | 123.89 |
| Oxytocin signaling pathway | 8/154 | 1.30E-04 | 9.16 | 110.15 |

**Table 3.**
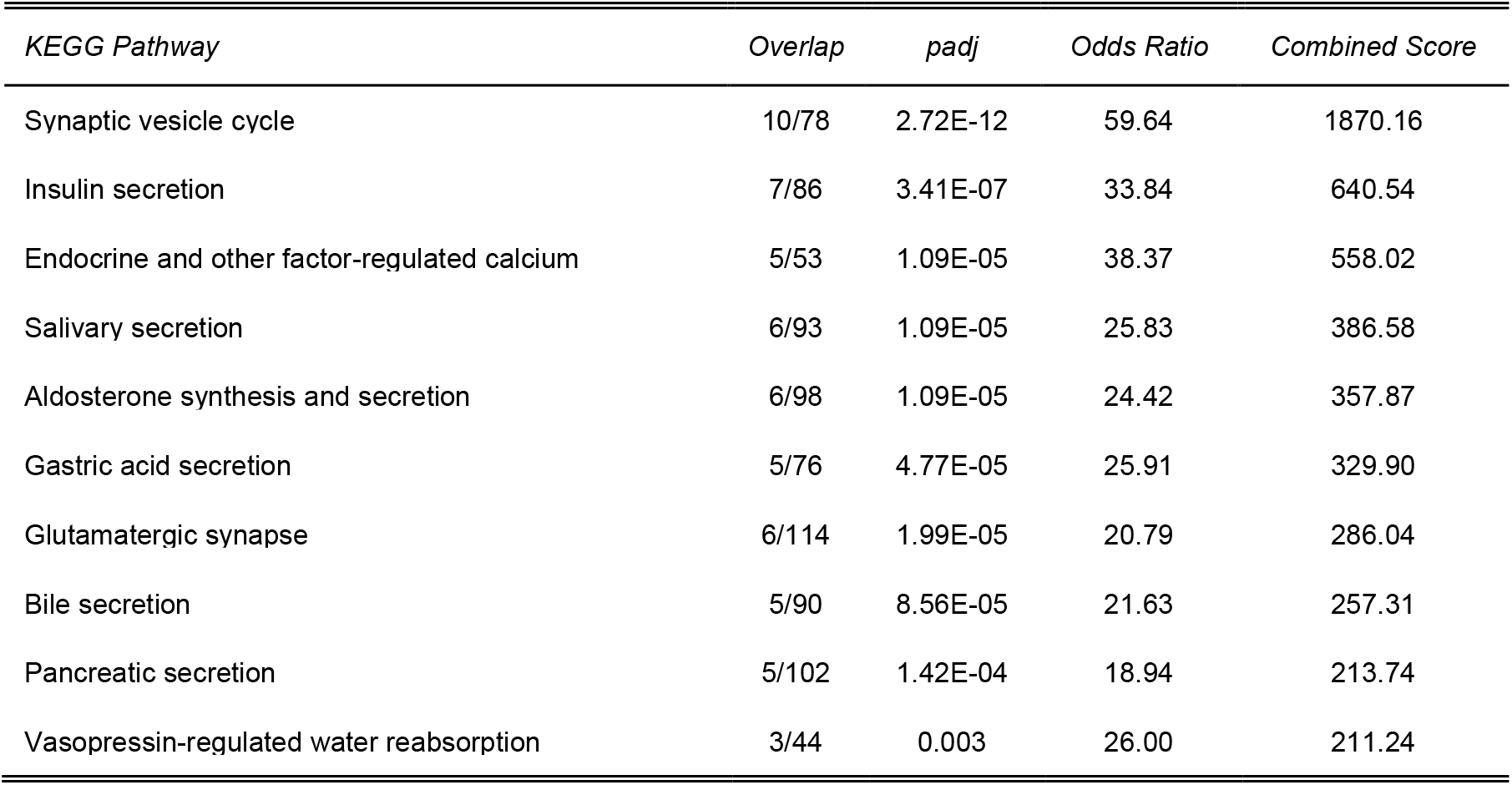
KEGG 2021 pathway enrichment for the synaptic vesicle and secretion co-expression module. (Module 2)

**Table 4.** Demographics of study groups.

|  | *Control* | *MDD* |
| --- | --- | --- |
| **Sample size** | N = 216 | N = 241 |
| **Age (years, avg)** | 47.66 (sd = 15.18) | 46.78 (sd) = 15.21) |
| **Gender (N, %female)** | 62 (28.70%) | 103 (42.74%) |
| **PMI (hours, avg)** | 24.08 (sd = 16.22) | 26.32 (sd) = 16.24) |

**Table 5.**
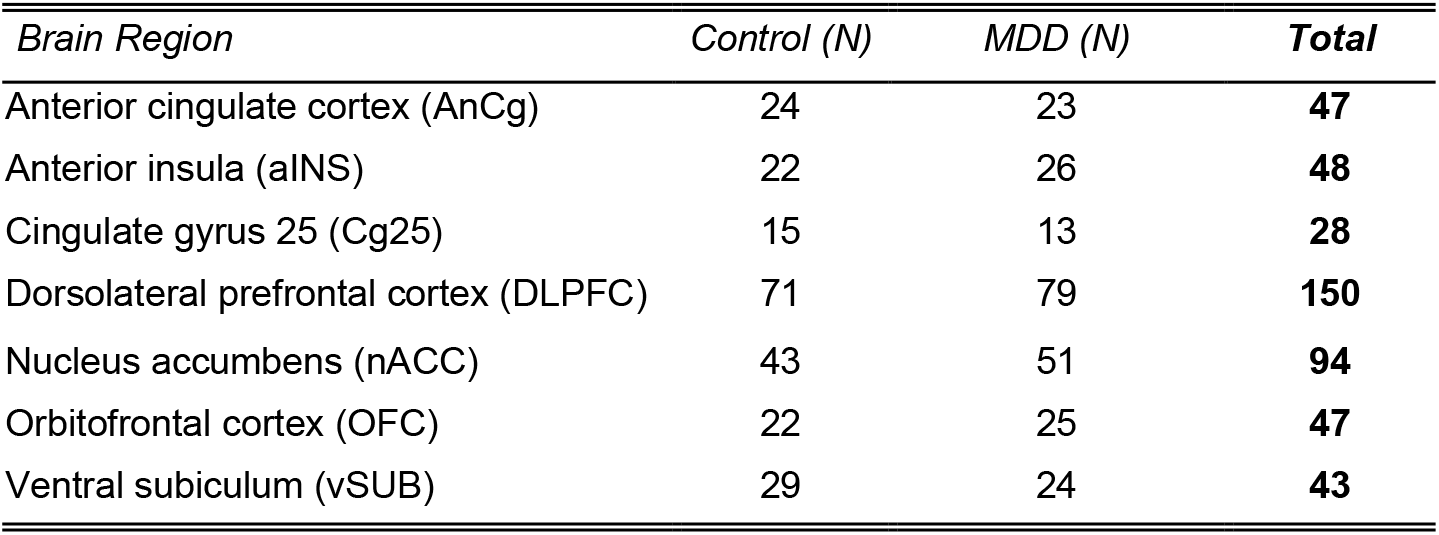
Distribution of samples by brain regions and the study groups that they belong to.

| *Brain Region* | *Control (N)* | *MDD (N)* | ***Total*** |
| --- | --- | --- | --- |
| Anterior cingulate cortex (AnCg) | 24 | 23 | **47** |
| Anterior insula (aINS) | 22 | 26 | **48** |
| Cingulate gyrus 25 (Cg25) | 15 | 13 | **28** |
| Dorsolateral prefrontal cortex (DLPFC) | 71 | 79 | **150** |
| Nucleus accumbens (nACC) | 43 | 51 | **94** |
| Orbitofrontal cortex (OFC) | 22 | 25 | **47** |
| Ventral subiculum (vSUB) | 29 | 24 | **43** |

We further investigated pathways enriched for individual co-expressed gene modules. We called the first module as “glutamatergic signaling module” because we observed a strong enrichment for the addiction (Gass & Olive, 2008; Tzschentke & Schmidt, 2003), glutamatergic synapse, and circadian entrainment (Biello et al., 2018; Chi-Castañeda & Ortega, 2018) pathways that were mainly controlled by AMPA (α-amino-3-hydroxy-5-methyl-4-isoxazolepropionic acid) and NMDA (N-methyl-D-aspartate) glutamate receptor activity. GRIN1, GRIN2B, and GRIA2 are the glutamate receptors that were identified in this module. Lastly, we called the second module “synaptic vesicle and secretion module” because it contained genes responsible for the transportation of ions such as Ca^2+^. Therefore, the synaptic vesicle cycle, different secretion-related pathways, and pathways related to calcium absorption are highly enriched. Negative enrichment scores of these two modules for the MDD suggest that glutamatergic signaling activity is downregulated for the brain regions where we observed NPAS4 as a common downregulated gene.

DGE and co-expression analyses are designed to identify linear associations in gene expression in pre-determined conditions (e.g., disease and control). Thus, in addition to DGE analyses, we took a machine learning-based approach called multiple-kernel learning (MKL) to identify non-linear associations between biological pathways and disease conditions. Previously, the same computational framework was applied to identify features that can predict the stages of cancer (Rahimi & Gönen, 2018) and the survival of individuals (Dereli et al., 2019). In our analysis, KEGG pathways were used to identify the informative gene groups to discriminate MDD patients from the control group. In this method, each pathway was mapped to a gene expression matrix, and distinct kernel matrices were calculated for each pathway. Using the optimized weighted combination of these kernel matrices, the algorithm finds a sparse set of pathways by discarding uninformative ones from the collection. We can infer the relative importance of the pathways by considering their resulting kernel weights. We used the normalized gene expression values from all brain regions and samples (n=457) to identify the common underlying biological mechanisms associated with MDD.

We reported the area under the receiver operating characteristic curve (AUC) values over 100 replications to evaluate the algorithm’s performance. The predictive performance of the MKL algorithm is increased when we included samples from all regions compared to three regions containing differentially expressed genes (Figure 3A) indicating that including more brain regions and samples in the analysis increases the reliability of the prediction model. We achieved an average AUC score of 0.83 with a standard deviation of 0.04 for the model including all brain regions (Figure 3A). 21 pathways were selected as informative, at least in 50 replications (Figure 3B). Pathways “Linoleic acid metabolism,” “Viral protein interaction with cytokine and cytokine receptor,” “Olfactory transduction,” “Staphylococcus aureus infection,” “Chemical carcinogenesis – DNA adducts,” and “Graft-versus-host disease” were selected as informative in all replicates. Because some of the chosen pathways do not directly relate to brain tissue, we would like to elaborate on the results by categorizing them based on the gene groups they share. Hence, understanding commonalities between these pathways would guide us better. Thus, we divided pathways into two main categories based on their functional relevance and gene composition. The first cluster contained eight pathways (Linoleic acid metabolism, Chemical carcinogenesis – DNA adducts, Ovarian steroidogenesis, Primary bile acid biosynthesis, Fat digestion and absorption, Maturity onset diabetes of the young, Metabolism of xenobiotics by cytochrome P450, Retinol Metabolism, and Drug metabolism – cytochrome P450) containing genes related to synthesis, absorption, and metabolism of lipids. In this group, genes related to the cytochrome p450 (CYP) family are abundant and shared between different pathways. Previous studies have focused on variants in CYP genes and their association with SSRI metabolism and the effectiveness of the treatment (Hodgson et al., 2013; Shalimova et al., 2021; Thakur et al., 2007; Veldic et al., 2019). On the other hand, our approach puts forward the idea that they can be used for diagnosis. “Nitrogen metabolism” and “Maturity onset diabetes of the young” can also fit in this category because they are related to metabolism. Several studies (Gu et al., 2021; Mocking et al., 2021) demonstrate the role of metabolism in patients with MDD. The second major group contained five pathways (Viral protein interaction with cytokine and cytokine receptor, Staphylococcus aureus infection, Graft-versus-host disease, Hematopoietic cell lineage, and Cytokine-cytokine receptor interaction) related to inflammation and immune system which is parallel to the enrichment of differentially expressed genes that we identified. The remaining four pathways were related to perceiving external stimuli through receptors (Olfactory transduction, Neuroactive ligand-receptor interaction, and Phototransduction) and glycosylation (Mucin type O-glycan biosynthesis). Overall, using KEGG pathways as features, we discriminated against MDD patients with high accuracy. The pathways we identified as discriminative can serve as a starting point for the research on MDD diagnosis.

**Figure 3:**
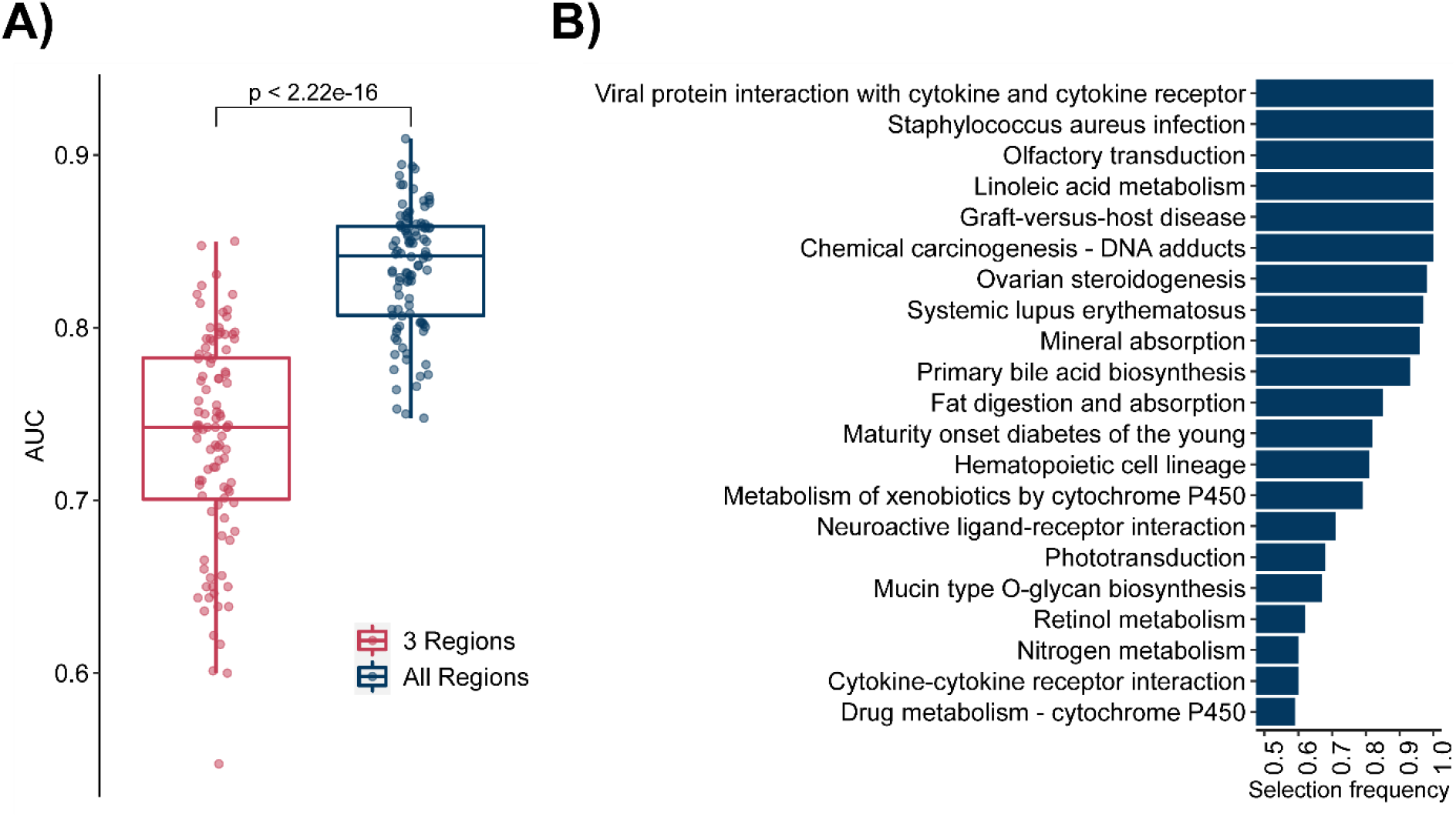
Multiple kernel learning results. A) Pathways selected as discriminative in 100 replication more than 50 times. B) Area under curve comparison of multiple kernel learning for 100 replications.

## Discussion

Our study combined multiple publicly available RNA-Seq datasets to identify novel pathways and genetic markers associated with MDD. A large sample size increased the sensitivity of the analysis, which led to the discovery of novel gene-disease associations. On the other hand, combining datasets from different sources introduces a certain amount of noise to the analysis. Moreover, Brodmann areas, individual segments of the cerebral cortex defining boundaries of each brain region, for a given region might slightly differ between studies. Therefore, we performed a preliminary quality filtration step to reduce noise and used the “study” covariate in our DGE analysis to eliminate some of that noise.

Our results show that the dorsolateral prefrontal cortex is the most affected region based on the number of differentially expressed genes, and downregulation of NPAS4 is observed for multiple brain regions. It should be highlighted that the larger change observed in DLPFC can also be attributed to its larger sample size. It has been previously demonstrated that NPAS4 plays a role in memory (Sun & Lin, 2016), modulating inhibitory-excitatory balance (Lin et al., 2008; Opsomer et al., 2020; Spiegel et al., 2014), epileptogenesis in mice, cocaine-induced hyperlocomotion (Lissek et al., 2021), cognitive well-being and many other diseases (Coutellier et al., 2012; Fu et al., 2020; Funahashi et al., 2019; Maya-Vetencourt, 2013). While the association between NPAS4 and MDD has been shown in mice previously (Jaehne et al., 2015), we validated the same relationship for humans and multiple brain regions. Supporting our findings, Gu et al. showed that patients with post-stroke depression had lower expression levels of NPAS4 in their peripheral blood mononuclear cells (Gu et al., 2019), which makes NPAS4 a potential diagnostic biomarker in the future. Our study suggests the central role of NPAS4 in major depression as an association factor. Although this study suggests a potential causation role of NPAS4 in the downregulation of synaptic plasticity in MDD, this hypothesis needs to be tested experimentally in model species.

To highlight the role of NPAS4 and understand that the relationship between differentially genes and co-expressed gene modules, we gathered our findings into an MDD model (**Figure 4**). In this model, we combined our findings with the existing literature on connections between genes and pathways. As we previously mentioned, combined differential gene expression analysis of DLPFC, nACC, and vSUB regions revealed downregulation observed within many immediate early genes (IEGs) such as NPAS4, FOS, FOSB, EGRs, NR4As, and ARC. It has been shown previously that FOS, FOSB, and their splice variants (Gajewski et al., 2016; Stone et al., 2008; Vialou et al., 2010) are associated with motivation and depressive behavior (Yi et al., 2019). Also, some antidepressants have been shown to increase the expression of FOS (Stanisavljević et al., 2019) and NPAS4 (Guidotti et al., 2012). These immediate early genes play important roles in maintaining essential synaptic functions (Gallo et al., 2018; Lanahan & Worley, 1998; Minatohara et al., 2016). It is known that the expression of IEGs is induced after neuronal stimulation, and NMDA glutamate receptors can induce the expression of IEGs through controlling Ca^2+^ influx (Greenberg et al., 1992; Xia et al., 1996) within neurons. ChIP-seq enhancer data of NPAS4 within mouse cortical neurons show that NPAS4 regulates immediate early genes (Kim et al., 2010). Differentially expressed genes and this experiment include IEGs in common; FOS, FOSB, NR4A1, NR4A3, JUNB, and NPAS4 itself. ChIP-seq data of NPAS4 embryonic mouse 14 days medial ganglionic eminence (mostly containing excitatory neurons) and cortex (mostly inhibitory neurons) shows that NPAS4 regulates distinct sets of late-response genes in inhibitory and excitatory neurons (Spiegel et al., 2014). In our DGE analysis, we observed a change in PTGS2, ATF3, ETV3, and CSRPN1 which were regulated in inhibitory neurons. By controlling the expression of other IEGs, which are also transcription factors, NPAS4 indirectly regulates the expression of many different genes as a master transcription factor. In our analysis, we observed a significant downregulation trend in these pathways that cumulatively lead downregulation of synaptic plasticity, circadian entrainment (Bunney et al., 2014; Lam, 2008; Walker et al., 2020), and learning abilities (long-term potentiation) in patients with MDD.

**Figure 4:**
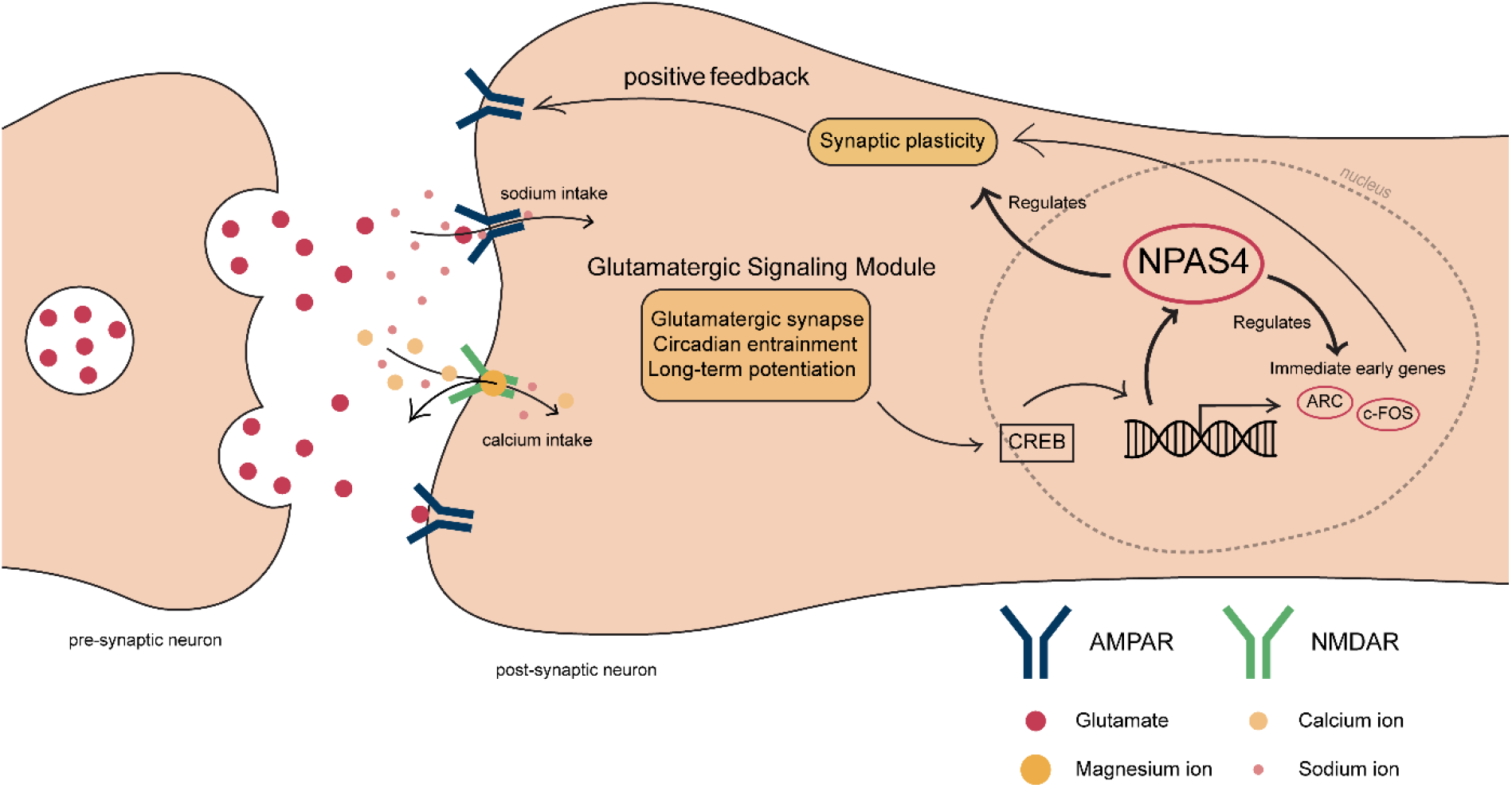
Summary of differentially and co-expressed genes and the enriched pathways

Current therapeutic strategies for MDD mainly target aminergic receptors such as serotonin and dopamine receptors (Harmer et al., 2017). These monoamine-oriented treatments have been ineffective, especially for patients with treatment-resistant depressions (Daly et al., 2018; Rush et al., 2006). Therefore, it is necessary to develop more effective therapeutics. This can only be achieved by understanding the molecular basis of the disorder. In this case, our study suggests that glutamatergic receptors can be used as drug targets in the treatment of MDD.In parallel to our findings NMDA antagonist ketamine and its enantiomer esketamine was shown to be effective for patients with treatment-resistant MDD (Daly et al., 2018; Dang et al., 2014; Ionescu et al., 2020; Canady et al., 2020). Although esketamine is an antagonist of the NMDA receptor, it leads to the activation of AMPA receptors (Sanacora & Schatzberg, 2014) that increase synaptic plasticity. Furthermore, the trial of NMDA co-agonist glycine induced the depressive state in mice (Salim et al., 2020). Thus, we conclude that the results of these drug trials are in line with the model we proposed in this study. We anticipate that antidepressants targeting glutamatergic signaling pathways will gain more popularity.

## Materials and Methods

### Datasets

In this study, three post-mortem RNA-seq datasets from Gene Expression Omnibus (GSE101521, GSE80655, and GSE102556) (Labonté et al., 2017; Pantazatos et al., 2017; Ramaker et al., 2017) were combined to increase the sample size and perform a statistically significant analysis of the MDD profile. A total of 216 control (28.70% female) and 241 major depressive disorder samples (42.74% female) were investigated based on their gene expression profiles. The average age of death of CTRL and MDD samples are 47.66 and 46.78, respectively. Samples from 7 brain regions, including the dorsolateral prefrontal cortex (DLPFC), nucleus accumbens (nACC), ventral subiculum (vSUB), anterior insula (aINS), anterior cingulate cortex (AnCg), cingulate gyrus 25 (Cg25), and orbitofrontal cortex (OFC) were analyzed.

### Data Analysis

#### Quality Trimming

FASTQC 0.11.7 (Andrews, 2010) was used to check the quality of each sample. We eliminated some of the samples directly from the analysis due to having very low quality in general. For the samples having low quality towards the 3’ end, we used Cutadapt (Martin, 2011) with the “--quality-cutoff 10” option. After performing 3’ trimming we concatenated fasta files for each patient when there are multiple fasta files for a single patient.

#### Alignment to the Human Genome

TopHat 2.1.1(Trapnell et al., 2009) was used for aligning reads to the human genome (GRCh37) (Church et al., 2011). At this step, we converted fasta files into bam files. Then by using the samtools (Li et al., 2009) sort option we converted bam files to sam.

#### Read Count

HTSeq (Anders et al., 2014) was used to obtain read counts for each patient. The distribution of counts for each region is given in Figure 1. Ensembl GRCh37 annotation list was used as a reference.

#### Differential gene expression analysis

Differential gene expression analysis based on the negative binomial distribution was performed in R with DESeq2 package (Love et al., 2014). Genes that significantly differentially expressed (adjusted p-value < 0.05) between major depressive patients and the control group were identified regarding sex, age, study and brain region that the sample is obtained from, and post-mortem interval covariates (full model, design ~ sampleGender + sampleAge + PMI + brainRegion + condition; brain region-specific model, design ~ sampleGender + sampleAge + PMI + condition).

#### Co-expression analysis

R package CEMiTool was used to perform co-expression analysis. Normalized count data from DLPFC, vSUB and nACC were included in the analysis. Variance stabilizing transformation was not applied before filtering the genes and default filtering p-value was used (0.1). Label of each sample was provided to obtain normalized enrichment scores for each of the modules in control group and MDD patients.

#### Identification of MDD-Associated Pathways Using MKL Algorithm

A multiple kernel learning (MKL)-based machine learning approach (Rahimi & Gönen, 2018) was used to identify informative pathways in discriminating MDD patients. Instead of first identifying the expressed genes and then performing a gene set enrichment analysis using these selected genes, the proposed MKL-based algorithm considers whole expression matrix and each pathway from the given collection at the same time. In this method, each pathway is mapped to a different kernel function using the expression profiles of the genes in the given pathway. Kernel functions are defined as the similarity measures between pairs of samples, and it is known that weighted combination of several kernel functions (i.e., MKL) increases the predictive ability of the kernel-based methods (Gönen & Alpaydın, 2011). At the end, the proposed method converges to a solution where kernels with non-zero weights are included in the final model for the classification. We considered that a pathway is selected to be used in the final model if the corresponding kernel weight was greater than 0.01.

The experimental setting that we used in machine learning model is as follows. We split our dataset by randomly picking 80% as training and 20% as test set. While splitting the data, we kept the ratio between the control group and MDD patients same in the training and test partitions. We repeated this procedure 100 times to obtain more robust performance measures and reported the experimental results over these 100 replications. We performed 4-fold inner cross-validation for selecting the model parameters (i.e., regularization parameter C). Since the gene expression is a count data, we first log2-transformed our dataset. Following that, we normalized the training set to have zero mean and unit standard deviation, while we normalized the test set using the mean and the standard deviation of the original training set. We followed the same computational setting as proposed in (Rahimi & Gönen, 2018) to obtain the relative importance of pathways.

#### Data and Materials Availability

The open-source code and supplementary data are available at our GitHub repository: https://github.com/CompGenomeLab/mdd-analysis

